# Role of surface negative charges in agonist binding to the ‘unliganded’ open state of the neuromuscular acetylcholine receptor

**DOI:** 10.1101/2024.12.23.630096

**Authors:** Nadira Khatoon, Tapan K. Nayak

## Abstract

Acetylcholine receptors (AChRs) expressed at the nerve muscle junction (NMJ) synapses are hetero-pentameric ligand-gated ion channels with two neurotransmitter binding sites (TBS) at α-δ and α-ε (adult)/γ (fetal) subunit interfaces. They are typical allosteric proteins which reside in at least two stable conformations: resting (**C**losed) and active (**O**pen) states. Mechanism of agonist (A) binding to the C state has been extensively studied, but agonist association to the O state has not been clearly understood. Here, by using engineered constitutively active AChRs, single-channel patch-clamp, and molecular dynamics (MD) simulations, we elucidate the differences between ‘unliganded’ (O) vs liganded (AO) active states and the mechanism of agonist association to the O state. Our results indicate that: 1. At the αγ-binding site, for a series of agonists, the O state agonist association rate (*j_on_*) was ∼30 times faster than the diffusion limit and agonist affinity was 2-orders of magnitude higher vs at the αδ- and αε-binding sites. 2. Electrostatic surface charge density owing to 4 negatively charged residues (γE57, γD113, γD174, and γE180) facilitates loop C capping, compaction of the TBS, and agonist stabilization in the AO state. 3. Mutating these residues in combination reversed the *j_on_* and binding affinity to those comparable to the αδ- and αε-binding sites. 4. φ-value analysis indicated the presence of a transition state intermediate between the O and AO states. Overall, we elucidate the role of neighboring charged residues outside the TBS in determining high-affinity agonist binding and their significance in shaping the synaptic response at the NMJ.

**Summary:** Nicotinic acetylcholine receptors (AChRs) are allosteric proteins that are crucial for muscle contraction. The isomerization of resting closed (C) to active open (O) state can be achieved by both bind-gate and gate-bind (rare) pathway. Little is known about agonist binding to the unliganded functional O state. Here, we elucidate the nature of unliganded O state and the mechanism of agonist binding to this high-affinity O state by single-channel current recordings, protein engineering, *in silico* methods and phi-analysis. Our results describe an energetic barrier between the unliganded and liganded O state owing to negatively charged amino acids near the agonist site. These charges partially contribute to the low- and high-affinity agonist energies specifically at the embryonic neurotransmitter binding site. We also discussed the physiological significance of these residues in shaping synaptic response.

**Significance Statement:** Embryonic-type AChRs are indispensable for nerve-muscle junction formation. Many mutations in these receptors cause congenital myasthenia syndrome by increasing constitutive channel activation. Therefore, understanding the nature of unliganded functional open (O) state is critical. Here, we describe the nature and the mechanism of agonist binding to the unliganded O (or apo) state. Single-channel currents were recorded with different background constructs and binding site mutations. The results highlight the importance of loop C and ligand orientation in determining high-affinity agonist binding. Further we elucidate the role of electrostatic interactions in agonist binding to the O state (high-affinity binding). We surmise that higher association rate of agonist binding to O state is the reason behind higher efficacy of the fetal receptor.

## Introduction

The acetylcholine receptors (AChRs) expressed at the nerve muscle junction (NMJ) synapses are typical heteropentameric ligand-gated ion channel receptors (LGICs) comprised of five homologous subunits; 2α_1_, β, δ, ε (adult)/γ (fetal) with neurotransmitter binding sites (TBS) located at the interfaces between α-δ and α-ε (adult)/α-γ (fetal). They are typical allosteric proteins that shuttle between at least two conformational states: resting closed (C) and active open (O), with O having higher affinity for the neurotransmitter vs the C state (1–4). AChRs rarely open in the absence of any agonist. In the presence of an agonist, the open probability of AChRs drastically increases (for example, by a million-fold due to acetylcholine (ACh)) owing to the higher affinity of O vs C.

A cyclic thermodynamic scheme best describes the mechanism of ligand binding and channel ‘gating’ in AChRs (1, 4). At the NMJ, most often the active functional O state of AChRs is attained via the bind-gate pathway (5), where ACh binds to the resting C state resulting in a low-affinity AC complex which isomerizes instantly to the high-affinity AO state. The resting unliganded receptor (C) may, however, rarely isomerize, but with a finite probability (∼*P_o_* = 10^-6^), to an unbound (apo), functional O state (6–8). The O state then binds to the neurotransmitter resulting in an active, agonist-bound AO state by the ‘gate-bind’ pathway (5). Many mutations which are implicated in congenital myasthenia syndrome (CMS) increase the probability of unliganded channel opening and, concomitantly, increase the progression through the gate-bind pathway (9, 10).

Formation of the low-affinity complex involves diffusion of the agonist from bulk solution, an encounter complex formation at the transition state, and a local conformational change (catch-hold-type ‘induced fit’(11)) or binding to a ‘primed’ C state (conformational selection (12)). However, the mechanism of high-affinity complex formation by agonist binding to O state is not clearly understood.

In recent years, the detailed structure of the neuromuscular AChRs has been elucidated in different functional states, including an α-bungarotoxin-inhibited non-conducting state (PDB ID: 6UWZ(13)), an apo closed and agonist-bound open structures embedded in lipid nanodisc (PDB Id:7QKO, 7QL5, 7QL6 (14)), a curare stabilized desensitized state (PDB Id: 7SMM; (15)) and resting (9AWK) and ACh bound (9AVU) state structures for fetal AChRs (16). The kinetic rates (*f_n_*, *b_n_*) and equilibrium dissociation constants (K_d_) for agonist binding to the low-affinity C state of the wild-type (WT) and several mutant AChRs have been measured in the past (17–23). The core binding pocket at both the TBS comprises of five conserved aromatic residues including αW149, αY190, αY198, αY93 and ε/γW55 or δW57 (24–26). Free energy changes associated with low-affinity agonist binding (ΔG_LA_) as a function of the different amino acid side chains, up to a resolution of molecular groups, have been experimentally quantified at all three types of binding sites for cholinergic agonists (αδ, αε, αγ) (23, 27, 28). Though the aromatic pocket is conserved at all three binding sites, they are asymmetric insofar as the ΔG_LA_ of the three TBS (ΔG_B_^αγ^ >> ΔG_B_^αε^ ∼ ΔG_B_^αδ^). Some of the residues within 20 Å of the core TBS (‘outer zone’ residues) are strongly implicated in modulating the C-state affinity and the asymmetry in binding sites (29).

On the flip side, the measurement of the kinetics and energetics of agonist binding to the unliganded O state has been challenging due to the (i) infinitesimally low probability of channel-gating in the absence of ligands; (ii) absence of appropriate concentration-response experiments in dramatically low concentration ranges (pM to nM) required for estimating the equilibrium dissociation constant for the high-affinity O state (J_d_); and (iii) the inherent complexity of unliganded gating (7, 8). Therefore, the mechanism of ligand binding to the active state is less clearly understood. Previously, there have been indirect attempts to measure J_d_ by assuming binding site symmetry and microscopic reversibility (30). Nayak et al., 2017 directly measured the high-affinity binding parameters for the αδ, αε and αγ sites from single-channel patch-clamp experiments and established microscopic reversibility as a rule in the thermodynamic cycle for some of the cholinergic agonists (5). However, ligand binding mechanism and ‘efficiency’ of ligand binding for structurally different classes of compounds are different in AChRs (31, 32). Therefore, high-affinity binding for non-cholinergic agonists may not be predictable from the cycle.

Further, molecular dynamic simulations (MD) have elucidated the structural changes associated with C vs O/D (13, 14) states. In LGICs, the low (AC) → high-affinity (AO) state transition involves asymmetric rotation and/or outward rigid body motions at the subunit interfaces at α_γ_-β-δ and γ-α_δ_ subunit interfaces (13–15). It has been observed in the carbamylcholine (CCh) and nicotine (Nic) bound O structure that several residues, including αK145, αD200, γE57, γN107, γV108, γD174, and γE180, change their positions in the principal and complementary subunits outside the core binding pocket (14). However, the structural dynamics involving C (resting) → O (apo), the nature of the apo O state (whether it is low or high-affinity state) and the local structural rearrangements between O → AO (high-affinity open) are not understood. Further, it will be interesting to know how these state transitions are facilitated by the residues outside the aromatic binding pocket.

Functional and structural dynamics studies so far to understand agonist binding to the O state suggest the following: (i) The position and orientation of the agonist head group were correlated with the high-affinity binding energy *in silico* (32) (ii) high-affinity binding is diffusion limited with the agonist bound receptor being O-like at the transition state (TS) (phi or ϕ ≈ 0; (5)) (iii) For some of the cholinergic agonists, *j_on_* at the αγ binding site is greater than the diffusion limit. Here, we hypothesize that the association rate greater than the diffusion limit owes to the negatively charged residues in the vicinity of the binding pocket and high-affinity binding proceeds through a transition pathway analogous to the low-affinity binding. We present the kinetics and energetics of ligand binding to the O state and investigate the role of the resident electrostatic charges outside the core binding pocket in determining the O state agonist affinity. Further, we illustrate the complex nature of the unliganded O state and the physiological significance of the gate → bind pathway. The results are discussed in the light of the structural dynamics studies performed based on the recent high-resolution cryoEM structures of AChRs.

## Results

### Thermodynamic cycle

Figure 1 illustrates the cyclic thermodynamic model for AChRs. In the absence of any agonist, C ↔ O gating transition of the receptor is governed by an equilibrium constant E_0_ (‘allosteric constant’; (7, 8, 33). The corresponding free energy change is the intrinsic gating energy (ΔG_0_). The equilibrium dissociation constants for C and O states are K_d_ (*k_off_/k_on_*) and J_d_ (*j_off_/j_on_*), respectively. The corresponding free energy changes associated with ligand binding to the C and O states are ΔG_LA_ (-0.59*ln(1/K_d_) at 23 ^0^C; LA: low-affinity) and ΔG_HA_ (-0.59*ln(1/J_d_); HA: high-affinity), respectively. Microscopic reversibility in this cycle has been elucidated for cholinergic agonists by rigorous measurements of the rates, equilibrium constants, and free energy changes by single ion channel current recordings (5). Due to microscopic reversibility in each half of the cycle ΔG_B_ = (ΔG_HA_ - ΔG_LA_) = ΔG_1_ - ΔG_0_.

**Figure 1.**
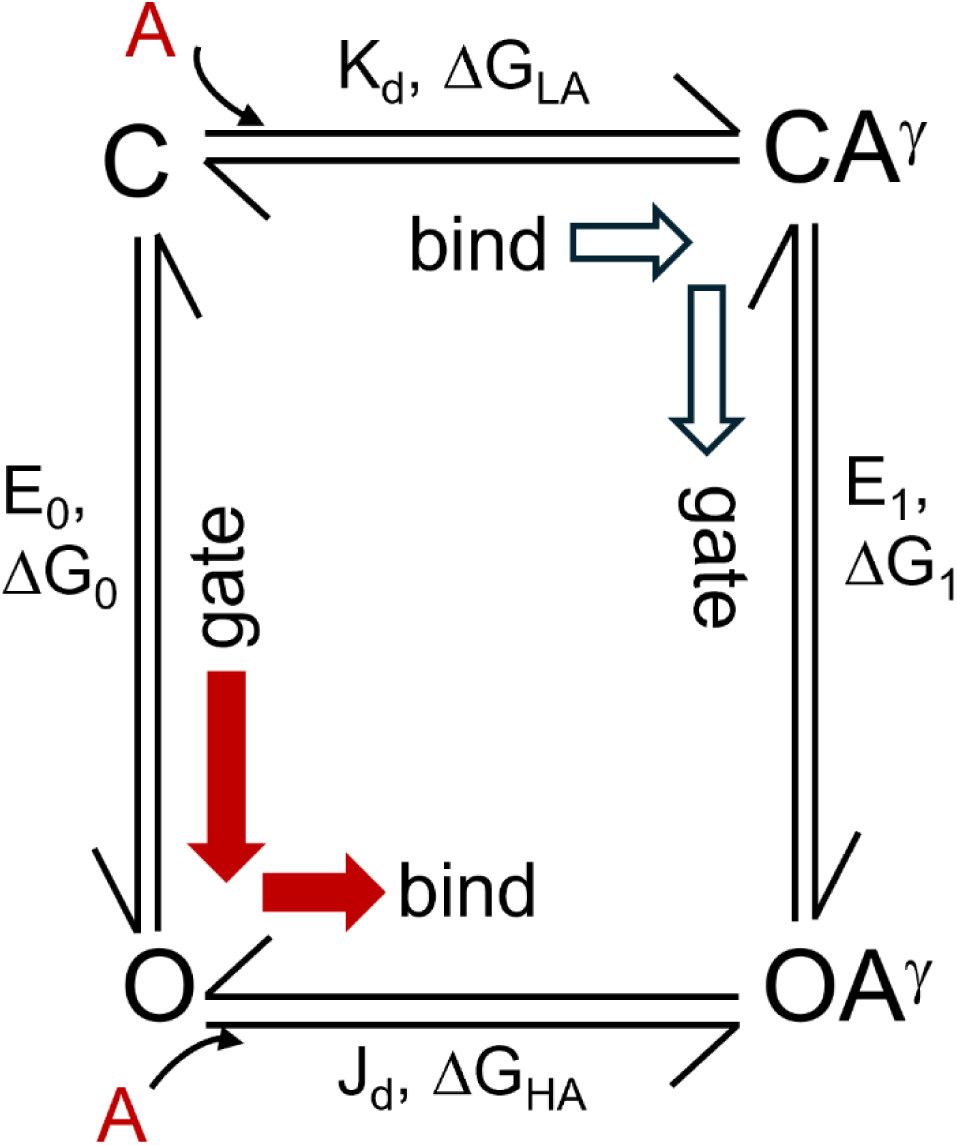
Cyclic model of AChRs. Thermodynamic cycle showing αγ-site (single-site) activation of AChRs. C (**C**losed), O (**O**pen), OA (**A**: Agonist) are the resting, active and agonist-bound active states. Vertical and horizontal arms represent gating and binding steps. E_n_ = gating equilibrium constants (n = 0,1 where n represents the number of bound agonists). ΔG_n_ = gating free energy. K_d_, J_d_ = low- and high-affinity dissociation constants. ΔG_LA_, ΔG_HA_ = low- and high-affinity binding free energies. The open and filled arrows show the two possible pathways for achieving stably bound open state (OA).

SI Figure 1 is the extension of the simple cyclic 1-site binding model, where binding to both the asymmetric binding sites (α-δ, α-γ/ε) are considered (cube model). The rate and equilibrium constants for the vertical and horizontal arms for ACh, CCh, tetramethylammonium (TMA), and choline (Cho) have been experimentally determined for the cube model (34) (SI Table 1). Here, we present the rate and equilibrium dissociation constants for the O state of AChRs in the presence of non-cholinergic agonists, Nic and anabasine (Ana).

### Unliganded or apo O functional state

Single-channel kinetic studies for the WT AChRs is challenging due to infrequent channel openings, hence, making studies involving agonist binding to the apo receptors practically impossible. Figure 2a elucidates the native WT adult and fetal AChR in the O state. To measure J_d_ of AChRs in the presence of Nic and Ana for the αγ-agonist site we knocked out the αδ-site and engineered high gain-of-function background mutations to ensure that the receptors are in the O state prior to agonist binding (see SI Methods). For instance, the background mutations βL262S, γL260Q, and δP123R in combination resulted in a gain-of-function of ∼1440000 (1.44×10^6^, (34)) folds, i.e. the ΔΔG_0_ for the combination was = -8.37 kcal/mol and ΔG_0_=1.68 kcal/mol (E_0_=0.058), which was comparable to the efficacy of the WT AChR in the presence of partial agonist Cho (E_2_=0.06). Figure 2b shows representative single-channel current records from three different combinations of background mutations of equivalent gain-of-function in the fetal AChRs. The single-channel current data were directly fitted by kinetic models as described in the SI Methods. The corresponding dwell-time distribution histograms were fitted with mixed exponential probability density functions (*pdfs*) (see SI Methods). Figure 2c shows the kinetic models that describe the unliganded single-channel currents for the three engineered background constructs. From the kinetic models, channel opening (*f_0_*) and closing (*b_0_*) rates were directly measured and gating equilibrium constant (E_0_) = *f_0_/b_0_*. Free energy change was estimated from the relation = ΔG_0_ = -0.59*ln (E_0_) (see SI Methods). The E_0_, ΔG_0_ and ΔΔG_0_ of the background combinations are given in SI Table 2. We chose βL262S+γL260Q+δP123R background for most of the J_d_ experiments in the fetal AChRs.

**Figure 2.**
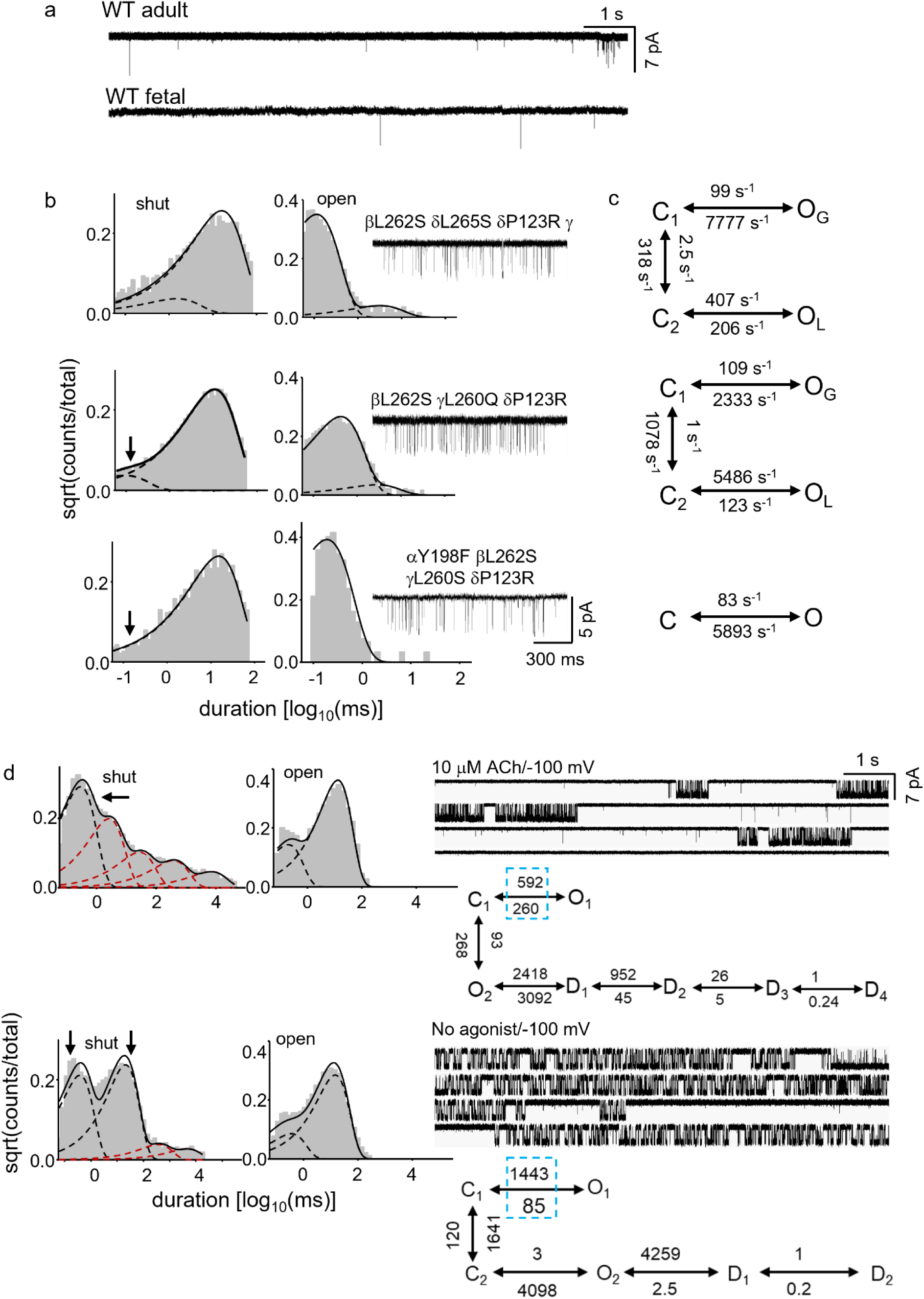
Comparison between unliganded O vs liganded OA. **a**. Low resolution single-channel current traces recorded from HEK 293 cells expressing WT adult and fetal AChRs. Note the low open probability of both the receptor types in the absence of agonists. **b**. Representative single-channel current records from AChRs having combination of background gain-of-function mutations (locations shown in figure) expressed in HEK 293 cells (right); Corresponding dwell-time duration distribution histograms are shown on the left. Solid spline lines: Sum of exponential probability density functions (*pdfs*); Dotted lines: component exponential *pdfs*. **c**. Kinetic model and the rate constants used to optimally fit the single-channel current interval data shown in b. **d**. (upper, right) Low resolution single-channel current data recorded from WT AChRs in the presence of 10 μM ACh (V_m_= -100 mV). The corresponding kinetic model that best fits the data is shown below. (lower, right) Low resolution single-channel current data recorded from AChRs with mutations αA96H and βV266A (-100 mV) in the absence of any agonists (V_m_= -100 mV). The corresponding kinetic model that best fits the data is shown below. The dotted blue box represents the gating rates obtained from the fit. The corresponding dwell-time distribution histograms are shown on the left. Solid spline lines: Sum of exponential *pdfs*; Red dotted lines: exponential *pdfs* representing the desensitization substates; Black dotted lines: exponential *pdfs* representing the gating state/substates (represented by arrow).

The kinetics of unliganded AChRs are inherently complex with multiple non-conducting [2 C and 3-4 D (Figure 2d)] and conducting (at least 2 O) states. However, in all the cases it was possible to identify the gating C and O components as the major components of the distribution. The minor unliganded gating component occupied ≤ 10% area of the exponential *pdf* and represented the long openings observed in unliganded channel activities. We refer to this minor component as O_L_ (L for long) and the major component as O_G_ (G for gating). In the presence of a mutation at one of the core binding site aromatic residues (αY198F, see Figure 2b; (5, 8)) the complex unliganded gating kinetics became relatively simpler as O_L_ was drastically reduced (gating can be fitted by simple C-O model). It has also been shown that the single-site knock out mutation (δP123R) and α-δ site specifically blocked by site-specific blocker, conotoxin (CTxMI) produce equivalent results in the case of cholinergic agonist (5). We propose that the minor gating component (O_L_) represents channel openings corresponding to a conformational state of the binding pocket which is akin to the high-affinity state and the receptor does not readily bind to agonists in this conformation (see below).

### Experimental determination of J_d_

In the gate-bind pathway, both O and AO states are functionally conductive states. To understand agonist binding to the O state, we first studied the kinetic differences between the O vs AO by comparing these states having approximately similar efficacy (ΔG) values achieved in the absence vs presence of agonist. We incorporated 2 high gain-of-function mutations αA96H and βV266A (34) in combination to attain ΔG_0_ values comparable to the liganded ΔG_2_ values in the presence of neurotransmitter ACh. Figure 2d shows the single-channel current traces and dwell-time duration distributions histograms for AChRs in the AO state (in the presence of 10 μM [ACh], upper) vs O state (with αA96H+βV266A, lower). In the presence of ACh, the closed duration distribution histogram showed only 1-major gating component and 4-D substates (Figure 2d), whereas the open duration distribution shows unbinding and 1-gating components. On the other hand, in the absence of any agonist, the C state duration distribution histogram shows at least 2-components: 1 major and 1 minor (see Figure 2b) and 2-3 D substates. The unliganded O state show relatively more complex activation kinetics vs AO. <u>We deal with the complexity of O vs AO elsewhere.</u>

To understand agonist binding to O, we focussed on the major gating component (O_G_). Figure 3a shows representative single-channel current traces, dwell-time duration distribution histograms in the presence of picomolar (pM) to nanomolar (nM) [Nic]. With increasing [Nic], an AO component progressively evolved (marked by arrow). The single-channel currents obtained with increasing [Nic] were globally fitted to the kinetic scheme shown in Figure 1 to obtain *j_on_* and *j_off_* (J_d_ =*j_off_*/*j_on_*). Figure 3a (right) shows the rates of agonist association and dissociation to the O state. We, then, measured the binding parameters for Ana, a tobacco alkaloid with similar chemical structure as Nic and an agonist of adult AChRs, to the O state of the fetal AChRs. Figure 3b elucidates the single-channel current traces, histograms and the kinetic model for the binding of Ana to the O state of the receptor. With progressive increase in the AO component (Figure 3b, marked by red arrow) with increasing [Ana], there was a simultaneous decrease in the major O component (marked by green arrow). This indicated that pre-existing unliganded O-state (O_G_) binds to agonists and manifests as the evolving AO component. The *j_on_*, *j_off_*, J_d_, and ΔG_HA_ for Nic and Ana are presented in SI Table 1. For both the agonists, the *j_on_* = 20.3×10^10^ M^-1^.s^-1^ and 18×10^10^M^-1^.s^-1^ respectively, which were more than 2-orders magnitude greater than the diffusion limit (1×10^9^). A value greater than the diffusion limit for the *j_on_* for Nic and Ana are consistent with the measurements for other cholinergic agonists (ACh, CCh, TMA and Cho; see SI Table 1). However, the average *j_on_* values for Nic and Ana were >6 fold greater than the average *j_on_* values for the cholinergic agonists. The *j_off_* for both the agonists are small and comparable to the cholinergic agonists (except Cho) indicating slower dissociation of the agonists from the O state.

**Figure 3.**
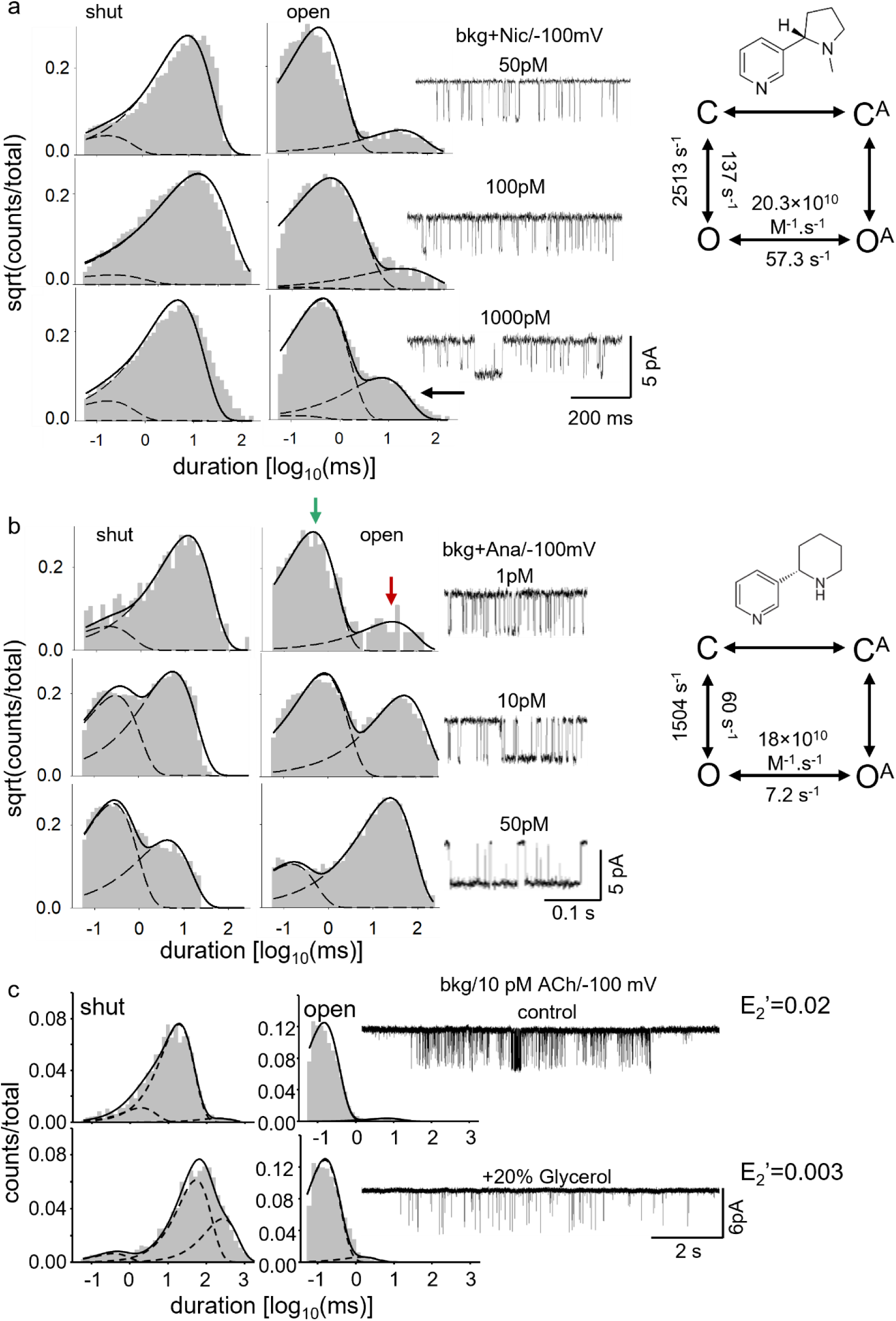
Measurement of J_d_ of AChRs in the presence of Nic and Ana. **a** and **b**. Representative single-channel current traces (right) and the corresponding dwell-time distribution histograms (left) in the presence of different concentrations of Nic (a) and Ana (b) (molecular structures shown). The C-O-AO kinetic model (right) was used to globally fit the single-channel current responses across concentrations. The kinetic rate constants obtained from the fitting are shown in the model. The horizontal arrow indicates the evolution of the AO (O^A^) component. Dotted lines are the exponential *pdfs* fitted to the current interval data for the traces shown on the right. The solid lines are the sum of exponentials fitted to the distribution. **c**. Representative single-channel current responses (right) and dwell-time distribution histograms (left) in the presence of 0.1 nM ACh (top) and 0.1 nM ACh + 20% glycerol (bottom). Bkg: Background (βL262S+δL265S+δP123R, see SI Table 1)

To further verify that the ligand association to the O state is diffusion limited and, perhaps faster, we performed binding experiments in the presence of pipette solution containing 20 % glycerol (viscosity =1.4 Pa.s as opposed to 0.01 Pa.s for water). We posited that in a viscous solvent medium the rate of diffusion would be altered, hence, the rate of agonist binding to the O state would proportionately decrease. Figure 3c shows single-channel currents and histograms elucidating ACh binding (0.1 nM ACh) to the O state of AChRs in the absence vs. presence of 20% glycerol. In the presence of glycerol, the channel *P_o_* was relatively less vs aqueous medium and the apparent E_2_ decreased by ∼7 fold. The corresponding rates and equilibrium constant value are given in SI Table 2.

### Elucidating agonist binding to the O state by all atom MD simulation

To understand why the *j_on_* rate constant for Nic, Ana and other cholinergic agonists is greater than the diffusion limit by more than 1-order of magnitude, we studied agonist binding to the TBS in the high-affinity state of AChR (see SI Methods). To that end, we performed all atom MD simulations (see SI Methods) using the CCh bound structures of AChR in the O state as the system (PDB ID 7QL6). Figure 4a (left) elucidates the electrostatic surface charge density superimposed on a space-filled model of the AChR in the apparent O state. The negative charge density in the O state structure was due to an array of negatively charged residues on the complementary side in the vicinity of the binding pocket including γE57, γD113, γD174, and γE180. Sequence analysis of the ε-(adult) vs γ-(fetal) subunits show that γE57 and γD113 are exclusively present in the γ-subunit which are replaced by Gly in the ε-subunit at the corresponding positions. Further, γD113 has been implicated in determining low-affinity agonist binding at the αγ-site. Additionally, γD174 was proposed to move closer to the binding pocket upon ACh binding (35). SI Figure 2a (left) elucidates the membrane view of the CCh-bound structure in the O state of AChR (7QL6.pdb (14)). The ECD is zoomed in to show the position of the above charged residues around the binding pocket (SI Figure 2a, right). The negative charge density seemed to manifest as a dipole moment along the subunit interface directed towards the neurotransmitter binding site (Figure 4a). We hypothesized that this electrostatic charge density in the O conformational state was critical in determining the high-affinity binding of the neurotransmitter to the TBS.

**Figure 4.**
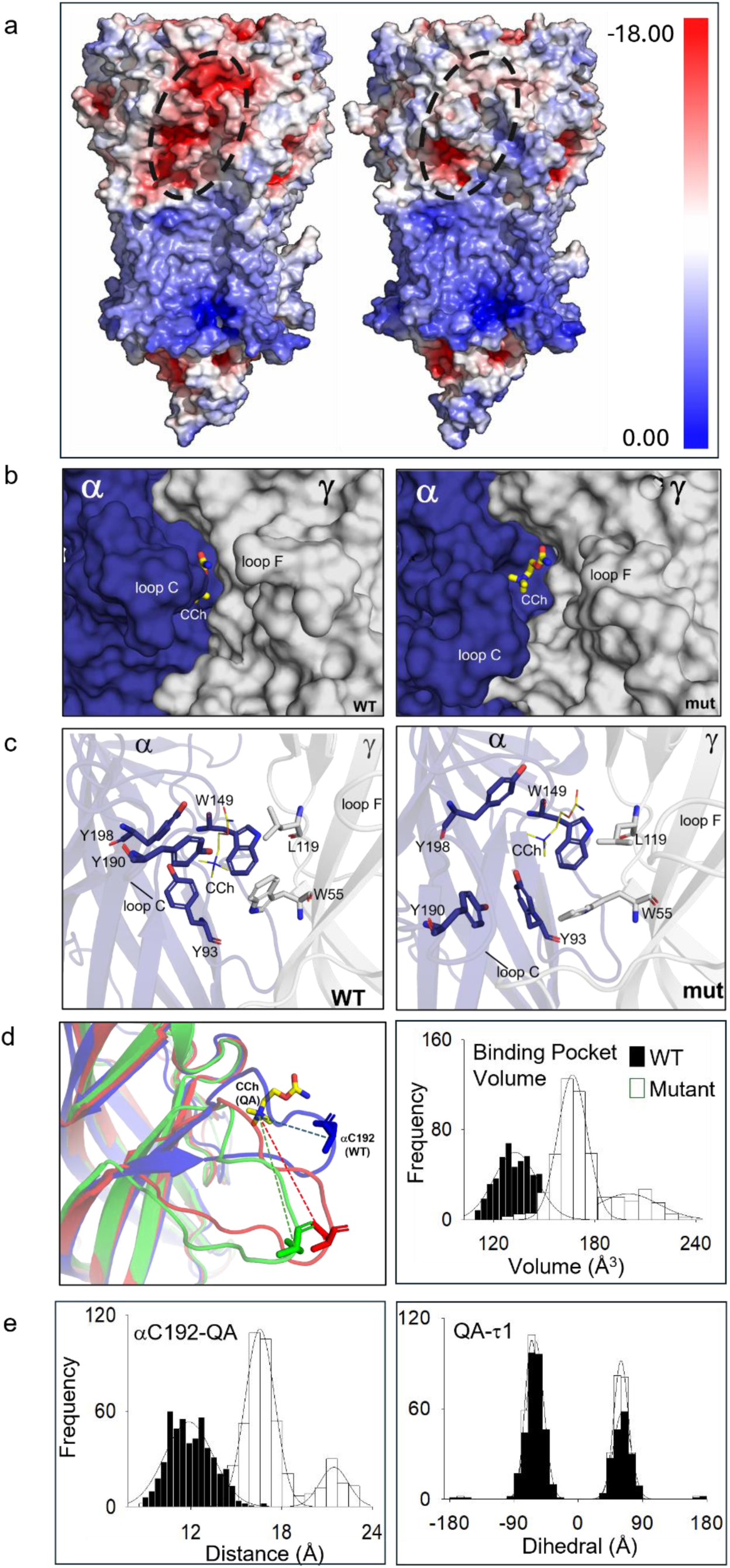
Binding site dynamics of the WT vs mutant AChRs. **a**. Electrostatic potential map of the αγ site of the WT (PDB ID: 7QL6) (left) and mutant receptor (right). Scale bar: Blue-positive, red-negative and white-neutral. Dotted circular region highlights the difference in surface charges between the WT and the mutant (or mut) receptor near the binding pocket. **b**. Surface representation of α-γ interface (Color coding: Blue-α, Grey-γ) highlighting CCh (yellow sticks) in the WT (left) and mutant (right) binding pocket. Note the difference in the sizes of the binding cavities of the WT vs mutant. **c**. Cartoon representation of the α-γ TBS interface showing the interacting residues within 4Å of the agonist (shown as sticks). **d**. (Left) Superimposed view of the TBS (represented as cartoons) of the WT vs 2 distinct conformations of the mutant captured during the simulation. Note the displacement of the loop C. The tip of the loop C (αC192) and CCh are shown as sticks and the distance between the two is shown as dotted lines. Color coding: Blue-WT; Red-mutant (major conformation, see Results); Green-mutant (minor conformation). (Right) Histogram showing the distribution of the volume of the binding pocket (αY93, αW149, αY190, and γW55) in the WT vs mutant receptors. Solid spline curve: Gaussian fit of the histogram peaks. Mean and standard deviation values of the fit are in SI Table 3. **e**. Histograms showing the distribution of the distance of αC192 residue (loop C) from the quaternary ammonium (QA) of CCh (left) and dihedral angle of CCh (τ1) (right). Spline curve: Gaussian fit of the histogram distributions. The mean and standard deviation values of the fit are in SI Table 3.

### In silico mutagenesis to understand high-affinity binding to O state

To understand the role of these charged residues, we mutated them *in silico* to Ala residues (Figure 4a, right). First, we mutated each of these residues separately and measured the charge densities in each case. SI Figure 3a shows the charge densities of AChRs in the presence of γE57A, γD113A, γD174A, and γE180A substitutions, respectively. There was no significant change in the charge densities compared to the WT receptor in the single mutants. Next, we mutated all 4 residues simultaneously to Ala. Interestingly, the charge density was drastically decreased in the receptor due to Ala substitution at these 4 positions (hereon will be called as the ‘mutant’). We measured the dipole moment for the ECD residues between 40-200 of chain A (alpha) and E (gamma) for the 7QL6 structure. The dipole moments for the WT and mutant were 1145D (Debye) and 873D, respectively (36).

### Conformational changes of the loop C

Capping movement of the loop C (displacement towards the aromatic binding pocket) has been implicated in the low-affinity binding of cholinergic agonists in the AChRs (22, 37) and more recently, shown during low (AC) → high (AO) affinity state transition (Singh et al, 2023). To understand the effect of the electrostatic charged residues *in silico*, we studied the conformational changes of the loop C for the WT vs the mutant receptors in our simulations. Figure 4c illustrates the residues interacting with the agonist within 4Å distance (αY190 is not within 4Å from the agonist in the mutant). The most conspicuous change between the WT and the mutant was the position of loop C. The average distance between loop C and B as well as loop C and the QA of the ligand increased in the mutant vs the WT. Figure 4d (left) shows the positions of loop C in the WT vs the mutant containing the quartet mutations. In the WT, the loop C was observed to closely appose against the binding pocket (‘capped’) for most part of the simulation, whereas in the quartet mutant receptor the loop C occupied conformations away from the agonist site (‘uncapped’). SI Figure 3b (left) is a histogram that elucidates the distance of the QA of the agonist from αY190 in the loop C that shows an increase between loop C and QA distance by 92±5% in the mutant (11.7±2.37 Å) vs the WT receptor (6.1±0.94 Å). Further, the average QA-αC192 (tip of the loop C, Figure 4e, left) and αW149-αC192 distances (SI Figure 3b, middle) increased by ∼42% and 34%, respectively, in the mutant vs the WT in the O conformation. In the quartet mutant, loop C preferably occupied 2 distinct positions around the binding pocket: a. completely uncapped (at 21.9±1.99 Å from the QA) and b. intermediate uncapped (16.4±1.33 Å from the QA) positions (Figure 4d, left). In the mutant, the loop C primarily occupied the intermediate uncapped conformation with >85% of the time during the simulations (Figure 4e, left). Overall, these results indicate that the loop C moves away or “uncaps” in response to the reduced negative surface charge density which might lead to weaker association and/or faster dissociation of the ligand. In the simulation of the high-affinity structure of neuromuscular AChR, loop F was found to close in onto the bound agonist leading to further stabilizing the loop C (14). In our simulations, however, the position of the loop F did not change significantly. However, the loop C-loop D distance in the mutant increased by ∼26% vs the WT. Apparently, the increase in this distance in mutant facilitates orientation of the non-α γW55 residue owing to repulsion of negative charges.

In concert with loop C positioning, in the mutant AChR, the TBS was loosely packed vs a compact binding site in the WT receptor in the O state (Figure 4d, right). Concomitantly, the neurotransmitter molecule was more snugly and deeply seated in the binding pocket in the O state of the WT vs the mutant AChRs (Figure 4b, c). Figure 4d (right) elucidates the variation in TBS pocket volume over the duration of the simulation (last 50 ns, see SI Methods). The volume of the TBS of the mutant receptor was ∼30% greater than the WT in the O state of the receptor (133.6±11.19 Å^3^ in WT vs. 173.1±18.59 Å^3^ in the mutant; Figure 4c). It may be further noted that the mutant binding pocket attains a bistable conformation as evidenced by 2 distinct volumes of the binding pocket (∼173.1 and 199.9 Å^3^, respectively, Figure 4d, right).

### Realignment of the agonist and the binding pocket

The position and orientation of the quaternary ammonium (QA) head group of the agonist in the binding pocket has been correlated with the O state binding energy (Tripathi et al., 2019). Therefore, we measured the orientation of the head group of the ligand in the TBS by measuring the N-C-C-O torsion angle (τ_1_). In the mutant receptor the QA head group was found to be bistable with τ_1_ = +62^0^ (∼50%) and -65^0^ (∼50%, Figure 4e, right) as opposed to a higher occurrence of a single conformation with counterclockwise torsion (τ_1_ = -62^0^ (>70%)) in the O state of the WT receptor. It indicates an increased fluctuation of the QA in the mutant receptor vs the WT.

In the fetal-type AChR γW55 is one of the special residues in the binding pocket that contributes a whopping ∼-5 kcal/mol binding energy in the presence of ACh in the low-affinity conformation (27). In the mutant, the favourable orthogonal positioning of γW55 with respect to the QA in the TBS is lost as it rotates by ∼100^0^ compared to the WT. SI Figure 3c is a distribution histogram for the side chain dihedral angle (χ1) of γW55 that shows rotation from +51° in the WT to -50° in the mutant.

Low-affinity agonist binding energy has been partially attributed to salt bridge and H-bond network in the vicinity of the binding pocket. We examined the salt bridge interaction between αK145-αD200 that weakened significantly in the mutant receptor. The average distance between αK145-αD200 increased from 3.1±1.1 Å to 4.8±0.8 Å (SI Figure 3d, left). This was further supported by H-bond occupancy data, which indicated reduced interaction strength in the mutant vs the WT (SI Figure 3d, right).

We then compared the α–δ binding interfaces of WT vs mutant to determine whether mutations in the γ-subunit influence the other binding site. The electrostatic potential map of the α–δ interface showed a similar charge distribution to the WT, with no significant alterations in the binding pocket. The average volumes of the binding pocket in the α–δ site of WT vs mutant were 139.7±4.09 Å^3^ and 137.7±4.45 Å^3^ (SI Figure 3e) and the salt bridge interaction between αK145-αD200 at α–δ site (average distance=3.2±1.06 Å) were comparable to those in the WT receptor. These results indicate that these mutations primarily affect the γ-subunit binding site, without altering the electrostatic environment or structural integrity of the α–δ interface.

### Effect of electrostatic charge neutralization on high-affinity binding

The C state affinity of the agonist has been shown to be modulated by some of the outer zone residues (outside 8 Å of the binding pocket), including γ/εD113 (29). For most of the charged residues the low-affinity binding kinetics have been studied and their K_d_ values in the presence of cholinergic agonists. The changes in ΔG_B_ values for γE57A, γD113A, γD174A, and γE180A residues were within ±0.6 kcal/mol (29). It indicates that the charged residues individually do not alter the low-affinity binding of the receptor.

We posited that though the charged residues do not alter the low-affinity binding, the high-affinity binding may be altered by the charged residues as indicated by the changes in the structure and charge densities in the O-state of the receptor. To study the effect of these charged residues in determining high-affinity binding we performed binding studies with the receptor having all 4 mutations in the presence of ACh as an agonist. We assume that the conclusions obtained for ACh binding is applicable to all cholinergic/non-cholinergic agonists as high *j_on_* values are observed for all of them. Figure 5a (top) shows the single-channel currents and dwell-time duration distribution histograms from the mutant AChRs with increasing concentrations of [ACh]. With increasing [ACh] a clear AO component evolved (marked by arrow) and the O component progressively declined. The dwell-time duration distribution histograms were fitted by mixed exponential *pdfs* obtained from the direct single-channel data fitting by the kinetic model shown in Figure 5a (bottom). The *j_on_* rate for ACh binding decreased by ∼10 fold and the *j_off_* increased by ∼30 fold in the mutant. The J_d_ of the mutant in the presence of ACh = 38 nM which was comparable to the αδ- and αε-sites (SI Table 4). The charge neutralization mutations in the vicinity of the binding pocket resulted in significant changes in J_d_ (380-fold) and ΔΔG_HA_ (+3.5 kcal/mol).

**Figure 5.**
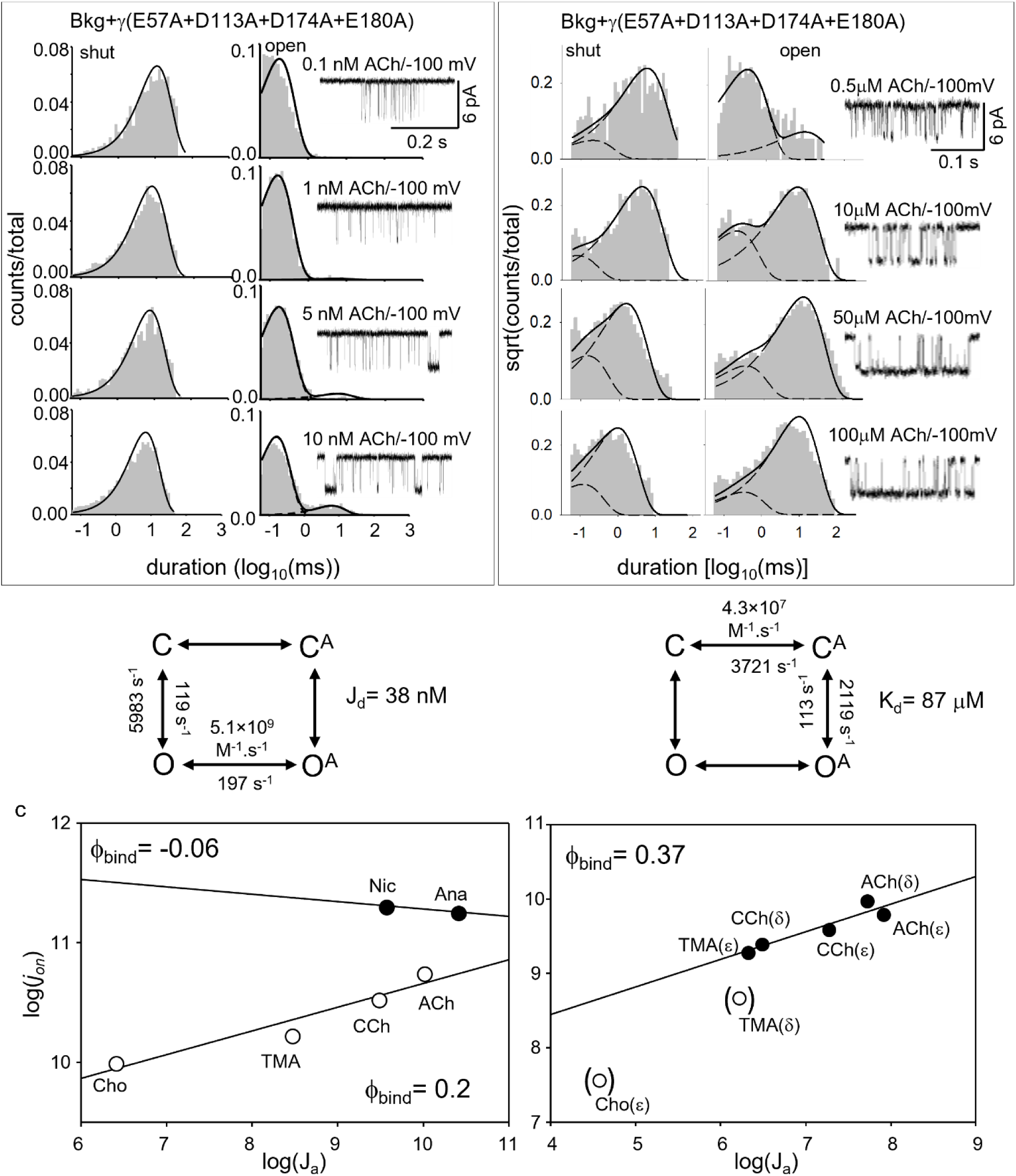
Low- and High-affinity binding parameters for AChRs with charge neutralization mutations. **a** and **b**. Representative single-channel current traces (right) and the corresponding dwell-time distribution histograms (left) with increasing [ACh]. The C-O-OA and C-CA-OA kinetic models (bottom) were used to globally fit the single-channel current responses for measuring gating/binding rate constants, J_d_ (a) and K_d_ (b). Dotted lines are the exponential *pdfs* obtained from the kinetic model fitting to the current interval data for the respective current traces. Solid lines: sum of exponentials fitted to the distributions. Bkg = βL262S+δL265S+δP123R. **c**. Scatter plot of the log (*j_on_*) vs log (J_a_) (a.k.a. binding REFER, see Results) for different agonists (ACh, CCh, TMA, Cho, Nic, and Ana) at the α-γ (left), α-δ and α-ε (right) sites. The straight lines are regression fits of the data passing through the scatter. ϕ_bind_ is the slope of the straight line. ϕ_bind_^α-γ^ = 0.2 (cholinergic agonists, R^2^= 0.92), -0.06 (Nic, Ana, R^2^= 1.0). ϕ_bind_^α-ε/δ^ = 0.37; R^2^= 0.88. Cho and TMA were unmasked outliers (shown by brackets).

To understand if the above mutations affect low-affinity binding, we recorded single-channel currents from AChRs with only functional αγ-site (αδ-site knocked out) in the presence of increasing μM [ACh] on appropriate gain-of-function backgrounds (see SI Methods). These background mutations favored transition along the bind-gate pathway like WT AChRs. Figure 5b (top) elucidates the single-channel currents, dwell-time duration distribution histograms and global fitting to the raw single-channel data by a simple 3-state (C-C_A_-O_A_) kinetic model (Figure 5b, bottom). The K_d_ of the mutant receptors = 86.5 μM as opposed to the K_d_ = 8.1 μM for the WT receptor. In the presence of mutations at the quartet of negatively charged residues, the low-affinity binding equilibrium constant increased by >10-fold whereas the high-affinity binding equilibrium constant increased by 386-fold. The E_1_* (at semi-saturated [ACh]) of the mutant combination as calculated from the fitting = 18.75. Hence, the ΔG_1_^mut^ for the mutant combination = -1.73 kcal/mol. Therefore, the ΔG_B_ of the αγ-site in the presence of the mutant combination = (ΔG_1_^mut^-ΔG_1_^background^) = -1.73-2.75 = -4.48 kcal/mol (see SI Table 2 for background information). On the other hand, K_d_/J_d_ for the mutant = 2249-fold and ΔG_B_= -4.55 kcal/mol. This indicates that the cycle is balanced in the case of the quartet mutations at the binding site.

### REFER analysis to study the transition state barrier

The relationship of rate and equilibrium constants for agonist binding can be understood by plotting rate-equilibrium constant free-energy relationship (REFERs) which provides insight into the relative temporal position of high-affinity binding event on an arbitrary time coordinate for ligand binding (2, 38). A REFER for agonist binding to the TBS was constructed by plotting the log change in J_a_ vs *j_on_* on the x- vs y-axis, respectively (Figure 5c). The slope of a REFER plot is phi (ϕ) that can have a fractional value between 1-0, where 1 represents the stable O, 0 represents the stable OA state and a fractional value represents intermediate conformations in between O and OA states. The ϕ-value for the WT high-affinity binding for cholinergic agonists = 0.2±0.02 and for Nic and Ana, the ϕ-value = ∼0. On the other hand, for the cholinergic agonists the ϕ-value = 0.37±0.01 at the αδ- and αε-binding sites. Further, we measured a 2-point REFER for the mutant from the measured binding rates. For the mutant AChR, the ϕ = 0.43.

## Discussion

The current work addressed some of the key questions pertaining to the O state agonist binding utilizing the embryonic AChRs as a model, such as: 1. what is the ‘nature’ and conformation of the unliganded O vs the liganded AO states? 2. Does agonist binding to the O state indeed barrierless and limited by diffusion? 3. What are the structural correlates which determine O↔AO transition? The results presented here elucidate the complex nature of the functional O state, suggest an intriguing possibility that agonist binding to the O state requires conformational selection between at least 2 pre-existing O substates (see below) and the surface negative charges in the vicinity of the TBS are critical in lowering the barrier between O and OA.

### Nature of the O state

The functional unliganded open state of the receptor for several gain-of-function mutations across the receptor (*Kampani et al, Unpublished*) shows at least 2 O substates, namely O_G_ and O_L_ (Figure 2c). Our results suggest that: 1. O_G_ represents the predominant gating substate, whereas O_L_ is a minor, relatively more stable open conformational substate. 2. In C↔O, the ϕ-value for the O_G_ > O_L_ substate indicating that at the barrier O_L_ is more open-like vs O_G_. 3. Of the 3 distinct conformations of the loop C, we propose that the tightly capped loop C conformation corresponds to the O_L_, whereas the uncapped, partially uncapped conformations (Figure 4d) correspond to O_G_. It is rational to visualize that a tightly capped loop C conformation will prevent an incoming agonist molecule whereas an uncapped loop C will accept an agonist and form a bound AO conformation. Agonists binding to only O_G_ was directly inferred from the observation that with increasing [agonists], there was progressive population shift from O_G_ to AO (Figure 3a, b). This is further corroborated by the following: 1. deletion or poly-Gly substitution in the loop C abrogates ligand dependent activation and O_L_, but major unliganded gating (O_G_) remained unaffected (39). 2. Mutations of the aromatic residues in binding pocket including at αW149, αY190, αY93 and αY198 (loop C and loop A residues) result in drastic reduction in long openings (8, 40). 3. αγ-site specific conotoxin, CTxMI binding leads to abrogation of the O_L_ component (5).

### Nature of the liganded AO state

The liganded AO state kinetics was more stable, less heterogeneous and had fewer number of C sub-states versus the O state. Further, the desensitization (comprised of 4-5 D substates) of the liganded AChR seemed different compared to the unliganded O AChR. This suggests that the two stable end states O and AO are 2 distinct conformations though they are both functionally capable of conducting ions. Further, in unliganded AChRs the number of ϕ-blocks observed were fewer than the liganded AChRs (*Kampani et al, Unpublished*), where ligand binding apparently resettles the structure in multiple phi-blocks. In a nutshell, a relatively more heterogeneous, flexible and complex O state becomes relatively homogeneous, ordered and simpler in kinetics in the presence of agonist.

### ‘Electrostatic steering’ and mechanism of agonist association to the O state

The quaternary ammonium (QA) head group (e.g. pyrrolidine group of Nic) of the cholinergic agonist interacts with 5 conserved aromatic residues (αW149, αY93, αY190, αY198 and γ/δ/εW55) (23, 41, 42) in the core binding pocket by a cation-pi interaction (21, 26, 43). The tail of the agonist is partially stabilized by γW118 and γL119 residues (14, 44). Though the core binding pocket residues are conserved, the binding sites at αδ, αε and αγ-subunit interfaces are functionally asymmetric (16, 34, 45). Several residues in the vicinity of the core binding pocket, including ε/γL104, γS111, γP112 and γD113, have been implicated in determining differential agonist affinity in (29). The 4 charged residues described here collectively contribute ∼ -1.4 and -3.55 kcal/mol to the low- and high-affinity agonist binding, respectively, enhance the high-affinity agonist association rate by 2-orders of magnitude higher than the diffusion limit and modulate low-affinity agonist binding. Increase in association rates beyond diffusion limit by ‘electrostatic steering’ (facilitatory effect of negative charges on a positively charged QA in agonist entry and low-affinity C state binding) has been observed in several enzymes including acetylcholine esterase (AChE; (46, 47), superoxide dismutase (SOD, (48)), and barstar (49–51). On the other hand, reduction of *j_on_* in the mutant to values similar to diffusion-limited reactions (∼10^9^ s^-1^) suggest that these vicinal negative charges (γE57, γD113, γD174, and γE180) strongly modulate high-affinity binding. It is, however, difficult to comprehend how the electrostatic effect is communicated to the core binding pocket though the conspicuous electric dipole moment is directed towards the TBS.

Further, the capped loop C conformation in the WT high-affinity OA state is facilitated by these negatively charged residues. Charge neutralization in the mutant AChRs resulted in bi-stable uncapped loop C conformation at the TBS. Further, the loop C capping was found to be weekly correlated with the ligand N-C-C-O dihedral (τ_1_). In the mutant receptor with neutralized charges, the QA head group was relatively more flexible and occupied 2 distinct conformations in the binding pocket indicating that these residues facilitate reorienting the positively charged QA in stable position (anticlockwise rotation) in the high-affinity O state. Recently, Singh et al proposed that AC → AO transition is associated with a ‘flip’ of the agonist (cis → trans reorientation about the long axis of the QA) and a ‘flop’ of the loop C (capping and uncapping) (52). In our simulations, we did not observe such flipping of the agonist though reorientation of the QA head group was observed. We propose that high-affinity agonist binding involves loop C capping and reorientation of the agonist in the binding pocket, which are both facilitated by electrostatic charges in the vicinity of the binding pocket.

### A distinct C↔AC barrier explains association rate greater than the diffusion limit

It has been proposed earlier (5) that at the α-δ and α-ε binding sites the high-affinity association rates (*j_on_*) for some of the cholinergic agonists were close to the diffusion limit in aqueous solution and the ϕ_bind_ for the high-affinity O state was estimated as ∼0. However, here we show that at the α-γ, TBS rearrangement significantly alters the position of the transition state for agonist binding (ϕ_bind_^WT^ = 0.2). In the mutant, neutralization of the charges further moves the position of this barrier to a later position with respect to the WT α-γ binding site (ϕ_bind_^Mutant^ = 0.4). Further, we estimated the ϕ_bind_ for the α-δ and α-ε sites as 0.37. Based on these observations we propose that there is a distinct barrier between O to AO in the WT AChRs which is lowered by the negatively charged residues at the αγ-site thereby resulting in *j_on_* rates which are 2-orders of magnitude greater than the diffusion limit. The reduction in the *j_on_* values (∼5×10^9^) in the case of the charge-neutralization mutant corroborate the conclusion drawn here.

### Physiological significance: Jd vs Kd

At the NMJ, typically, the neurotransmitter binds to the WT AChRs in the C state and follows the bind-gate pathway. Hence, K_d_ and efficacy (E_2_) determines the activation kinetics of the synaptic currents at the NMJ. However, in AChRs, several gain-of-function mutations have been characterized (9, 10) including those which cause congenital myasthenia syndrome (CMS) which result in increased constitutive channel openings. In such cases, high-affinity O state binding followed by gating becomes significant. To understand the role of J_d_ vs K_d_ in shaping synaptic response, we simulated macroscopic current response utilizing the cube model shown in SI Figure 1. Figure 6a elucidates the macroscopic currents generated at the synapses expressing WT, and a hypothetical mutant AChR with 10-fold increased J_d_. The difference in macroscopic response was significantly different at 10 μM [ACh]. We then compared the macroscopic current responses obtained from the WT and hypothetical mutations that increase K_d_ (K_d_ mutant) and J_d_ (J_d_ mutant) by 10-fold, respectively in the presence of 10 μM ACh (see SI Methods). In both the mutants the peak macroscopic currents decreased substantially though to different extents and the activation kinetics were significantly different from the WT (Figure 6b). In the case of the J_d_ mutation, the reduction in the macroscopic currents was more profound. Further, we measured concentration response (CRC) of WT and mutant AChRs in the presence of increasing [ACh]. Figure 6c shows the normalized and non-normalized CRC for the WT, K_d_ and J_d_ mutants. In the presence of the J_d_ mutants, the peak inward currents were significantly less than the K_d_ mutant. This indicates that the dissociation constant from the high-affinity O state determines the macroscopic currents to a greater degree vs similar perturbation to the low-affinity equilibrium dissociation constant. We surmise that in the fetal AChR the higher efficacy vs the adult AChRs may be attributed partly to the extremely low value of the J_d_.

**Figure 6.**
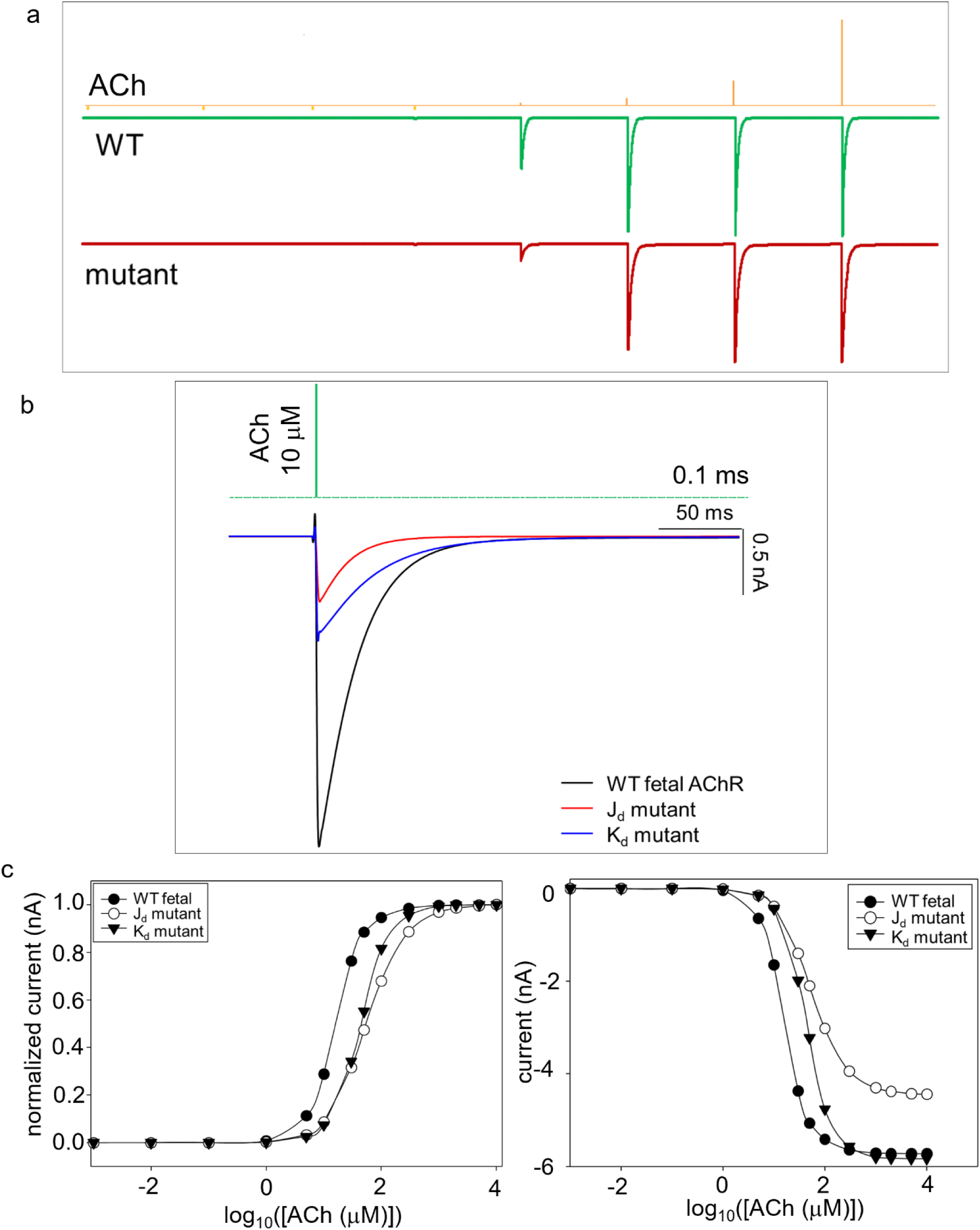
Simulated mEPCs at the developing NMJ. **a**. Simulated train of mEPCs generated using the model shown in SI Figure 1 and rate constants in SI Table 1 (pulse duration = 100 μs) for the WT and a hypothetical mutant AChR with 10-fold increased J_d_. [ACh] ranges from 1 nM to 10 mM. The timing of ACh pulse application is shown above the response (yellow trace). Significant AChR current responses were observed above >1μM [ACh]. **b**. Comparison of the macroscopic current responses obtained from the WT and model AChRs carrying hypothetical mutations that increase K_d_ (K_d_ mutant) and J_d_ (J_d_ mutant) by 10-fold, respectively in the presence of 10 μM ACh (black = WT; red = J_d_ mutant; blue = K_d_ mutant). **c**. Scatter plot showing the normalized (left) and peak (right) current responses in the presence of increasing [ACh]. The J_d_ mutant (open circle) and K_d_ mutant (downward filled triangle) are right shifted with respect to the WT (filled circle) AChRs. The solid lines are sigmoidal curve fitting to the scatter plot.

## Methods

We performed cell-attached patch-clamp electrophysiology, and molecular dynamics simulations using mouse AChRs. Single-channel currents were analysed using QuB software to estimate rate constants, by fitting a kinetic model to intra cluster interval durations. A detailed description of the methods used is given in *SI Appendix*.

## Acknowledgements

NK is grateful to IIT Delhi for the research fellowship. The authors acknowledge IIT Delhi HPC facility for computational resources. The research was funded by SERB CRG (CRG/2022/007550), DBT (BT/PR47726/CMD/150/26/2023) and ICMR (IIRP-2023-0990) grant to TKN.

