## Supplementary Information for "Role of surface negative charges in agonist binding to the ‘unliganded’ open state of the neuromuscular acetylcholine receptor"

**Running Title:** Mechanism of Agonist binding to the unliganded open state of AChRs

**Authors**

Nadira Khatoon[§], Tapan K. Nayak[§]*

**Author affiliations**

[§]Kusuma School of Biological Sciences, Indian Institute of Technology Delhi, Hauz Khas, New Delhi-110016

***Corresponding Author**

Tapan K. Nayak,
Kusuma School of Biological Sciences
Indian Institute of Technology Delhi,
Hauz Khas,
New Delhi-110016


**Author contribution**

Conceptualization: TKN; Execution of research, data analyses, and interpretation, review of manuscript: NK, and TKN; Original manuscript preparation: TKN, NK

**Competing interest Statement:** The authors declare no competing interests.

**Key words:** Association rate, Electrostatic steering, Diffusion limit, single-channel, patch-clamp, molecular simulations

### SI Methods

#### Cell culture and transfection

Human embryonic kidney (HEK) 293 cells were maintained at 37°C (5% $CO_2$) in Dulbecco's minimal essential medium (DMEM) supplemented with 10% (v/v) fetal bovine serum and 1% (v/v) penicillin-streptomycin (pH 7.4). Cells were transiently transfected using calcium phosphate precipitation method (1) by incubating them with cDNAs of the subunits of muscle-type AChR (total 2.5 μg per 35 mm culture dish) in a ratio of 2:1:1:1 (α:β:δ:γ/ε). Most electrophysiological experiments were done ~24 hrs post-transfection.

#### Protein Engineering

Wild-type (WT) AChRs rarely open without agonists. Therefore, $J_d$ cannot be measured for the WT receptors directly. We engineered AChRs harboring gain-of-function mutations to increase the probability of constitutive channel activation and agonist binding to the unliganded/apo receptors. Mutations were incorporated using site-directed mutagenesis kit (NEB, MA, US) and were confirmed by nucleotide sequencing. The primers for mutagenesis were designed using the NEBaseChanger tool (NEB). We incorporated mutations away from the binding pocket and widely separated from each other to ensure minimal interference in agonist binding and reduce any potential inter-residue coupling (2). In neuromuscular AChRs, residues separated by >16 Å behave functionally independent of each other (3).

To experimentally measure the high-affinity equilibrium constant ($J_d$) we chose the gain-of-function background mutations (βL262S+γL260Q) that increase constitutive channel opening. These two mutations were far away from the TBS in the transmembrane domain of the β and γ subunits. Due to increased constitutive channel opening, the probability of AChRs following the gate-bind pathway and agonist binding to the unliganded conducting state of the channel were expected to increase. We further incorporated δP123R mutation to disrupt (knockout) the α-δ agonist binding site that allowed measurement of the rates for agonist binding (Nic and Ana) to the α-γ site only (4).

To investigate the role of electrostatic charges in regulating the $J_d$ of the fetal receptor, we identified all the charged residues within 25 Å of the binding pocket. Of these, we mutated 4 strategically placed negatively charged residues (γE57, γD113, γD174, γE180) to alanine (Ala). To measure the rates associated with high-affinity binding, we used a different background gain-of-function mutations in combination (βL262S + δL265S) to increase the constitutive activity of the receptor as described above. The kinetic parameters of the background mutations are given in SI Table 1.

**Electrophysiology**

Single-channel currents from AChRs were recorded in cell-attached patch configuration. The cells were bathed in $K^+$ ringer composed of 142 mM KCl, 5.4 mM NaCl, 1.8 mM $CaCl_2$, 1.7 mM $MgCl_2$, and 10 mM Hepes/KOH (pH 7.4). The patch pipettes were filled with Dulbecco's PBS containing 137 mM NaCl, 0.9 mM $CaCl_2$, 2.7 mM KCl, 1.5 mM $KH_2PO_4$, 0.5 mM $MgCl_2$, and 8.1 mM $Na_2HPO_4$ (pH 7.3/NaOH). Agonists of different concentrations were added to the pipette for liganded experiments.

Patch pipettes were fabricated from borosilicate glass, coated with sylgard (Dow Corning), and fire polished to a resistance of ~10-12 MΩ when filled with pipette solution. Single-channel currents were recorded using dPatch low-noise digital amplifier (Sutter Instruments, Novato, CA) at 10 kHz and digitized at a sampling frequency of 50 kHz.

**Kinetic Analysis**

Kinetic analyses of single-channel currents were performed by using the QuB software suite. To estimate rate constants, clusters of single-channel activity, flanked by ≥20 ms nonconducting periods (representing desensitization), were selected by eye. The clusters were idealized into noise-free intervals using the segmental K means (SKM) algorithm after digitally filtering the data at 10 kHz. The gating rate constants ($f_n$ and $b_n$ for forward and backward reactions) were determined from the inverse of the time constant ($\tau$) associated with the predominant ($\geq$80%) briefest component of each dwell time distribution histograms that are obtained by fitting the idealized data to kinetic models using maximum interval likelihood (MIL) algorithm. The gating equilibrium constant was calculated from the ratio of the rate constants, $E_n = f_n/b_n$. The total free energy change of the gating isomerization (kcal/mol) is $\Delta G_n = -0.59 \times \ln(E_n)$.

The equilibrium dissociation constants for the O state were estimated by globally fitting the single channel current data obtained at increasing agonist concentrations of ACh, Nic, Ana using the following kinetic scheme:

$$C \longleftrightarrow O^* \overset{A}{\longleftrightarrow} AO,$$

where, C is the resting state, O* is an active state, and AO is an agonist-bound open state. $J_d$ was estimated from the second step of the kinetic scheme from the measured ratio of $j_{off}$ and $j_{on}$. Since low concentrations of agonists were used, there was no channel block; thus, membrane voltage was kept at -100mV during all the experiments.

**Simulating Synaptic Currents**

The dose-response simulation tool of the QuB software suite was used to simulate synaptic currents. Miniature endplate currents (mEPCs) were simulated by using 1000 channels at different ACh concentrations (1nM to 10 mM) using pulse duration of 100 μs and sweep duration of 100 ms. The model used for simulating synaptic current is shown in SI Figure S1, and the kinetic rate constants are given in SI Table 1. The integrated current response was obtained by integrating the current over every 1 s time interval.

Since the energy of the AChR activation is conserved throughout the cycle,

$$\frac{E_2}{E_0} = \left(\frac{K_d}{J_d}\right)^2$$

From the above equation, $E_2 \propto K_d$, thus a 10-fold increase in $K_d$ would increase $E_2$ by 100, and $E_2 \propto 1/J_d$, increase in $J_d$ would decrease $E_2$ by 100 folds. We then compared the total synaptic response among WT-fetal, $K_d$ mutant, and $J_d$ mutant and plotted the results.

***In silico* Mutagenesis and Molecular Dynamics (MD) Simulation**

We used the recently solved carbamoylcholine (CCh) bound structure of *Torpedo californica* (PDB Id:7QL6) (5) for the present study. Charged residues near the binding pocket (γE57, γD113, γD174, γE180) were mutated to Ala using the Structure Editing module of UCSF Chimera v1.16 (6). A total of six simulation systems were prepared: a WT receptor system, four with single binding site mutations, and a system with all four mutations mentioned previously. The Dunbrack 2010 rotamer library was used to select the orientation of the mutated residue with the least atomic clashes. Before proceeding with simulations, the energy of the systems was minimized by 1000 steps of steepest descent with the step size of 0.02 Å.

The individual WT and mutated proteins were then embedded in a bilayer membrane that consists of 1-palmitoyl-2-oleoyl-sn-glycero-3-phosphocholine (POPC) of size 150Åx150Å using the Membrane builder plugin of VMD v1.9.4 (7). The membrane-embedded systems were then solvated with TIP3P water and ionized by adding 150 mM NaCl. The lipids and water in the vicinity of the membrane-embedded protein (within 1.5Å) were removed. The MD simulation was performed using NAMD v2.14 (8) with CHARMM36m as the force field parameter (9).

Initially, all the prepared systems were subjected to 50000 steps of energy minimization via the steepest descent method, followed by an equilibration of 20 ns by gradually reducing the harmonic restraints applied on the lipid molecules to avoid sudden deformation of the bilayer. Planar restraints were applied on the head group atoms of the lipid molecules, whereas dihedral restraints were applied on the unsaturated carbon atoms of the oleyl tails of lipid molecules. Subsequently, all the systems were simulated without any restraints for about 100 ns (varying with systems) under NPT conditions using standard periodic boundary conditions and minimum image convention. Periodic conditions were applied, and the Langevin dynamics method was used to set the temperature at 310 K and pressure at 1 atm. Using the SHAKE algorithm, an integration timestep of 2 fs was used while keeping all the covalent bonds involving hydrogen atoms constrained to their equilibrium bond length (10). The particle mesh Ewald algorithm was used to calculate electrostatic interactions (11). A switching distance of 10Å and a cutoff distance of 12Å were applied for non-bonded interactions. The simulations were run at a timestep of 2 fs, and the coordinates were saved every 100 ps for later analyses. All other parameters were left to the default values. The stability of the MD simulations was confirmed by plotting the root mean square deviation (RMSD) of the protein over the trajectory (SI Figure S2 b).

**Structural Analysis**

PDB2PQR v3.5.2 was used to calculate the charges on the protein at physiological pH. CHARMM force field was used since the simulations were run using the same (https://server.poissonboltzmann.org/pdb2pqr). We have used the Adaptive Poisson-Boltzmann solver (ABPS) to calculate the surface potentials of the representative structures (https://server.poissonboltzmann.org/apbs). ABPS uses the output generated from PDB2PQR as an input to describe the surface potential of protein by solving the Poisson-Boltzmann equation (PBE) (12). All the parameters for electrostatic potential calculation were kept at default values. We have used PyMol v2.5.7 to visualize and extract images (13). The system stability was analysed using RMSD and RMSF plots. All structural analyses were done from the last 50 ns of the production run (distances, dihedrals, and salt bridge) using VMD v1.9.4 (14). To identify H-bonds between the donor-acceptor residues, a cutoff of <3.5Å was used.

The shoelace calculation method determined the volume of agonist binding pocket (15). The graphing software SigmaPlot 11.0 was used to represent the data (16).

SI Table 1. Kinetic rates and equilibrium constants for all the binding sites of neuromuscular AChRs with different agonists.

| Site | Agonist | $j_{on}$ | $j_{off}$ | $J_d$ (nM) | $k_{on}$ | $k_{off}$ | $K_d$ (μM) |
|---|---|---|---|---|---|---|---|
| α-γ | ACh | $5.6 \times 10^{10}$ | 6 | 0.10 | $2.2 \times 10^{8}$ | 1794 | 8.15 |
| | CCh | $3.4 \times 10^{10}$ | 11.4 | 0.34 | $1.5 \times 10^{8}$ | 4335 | 28.7 |
| | TMA | $1.7 \times 10^{10}$ | 59 | 3.5 | $1.38 \times 10^{7}$ | 2672 | 194 |
| | Cho | $1 \times 10^{10}$ | 4041 | 404 | $5.8 \times 10^{6}$ | 5410 | 933 |
| | Nic | $8.6\times10^{10}$ | 57.3 | 0.67 | - | - | - |
| | Ana | $1.8\times10^{11}$ | 7.2 | 0.04 | - | - | - |
| α-ε | ACh | $6.3 \times 10^{9}$ | 78 | 12.4 | $1.18 \times 10^{8}$ | 18246 | 153 |
| | CCh | $3.9 \times 10^{9}$ | 211 | 54.1 | $7.2 \times 10^{7}$ | 13072 | 182 |
| | TMA | $1.94 \times 10^{9}$ | 933 | 481 | $1.78 \times 10^{7}$ | 14240 | 802 |
| | Cho | $3.7 \times 10^{7}$ | 1013 | 27200 | $1.52 \times 10^{6}$ | 10319 | 6788 |
| α-δ | ACh | $9.52 \times 10^{9}$ | 182 | 19.2 | $1.61 \times 10^{8}$ | 17136 | 106 |
| | CCh | $2.46 \times 10^{9}$ | 816 | 331 | $2.52 \times 10^{7}$ | 13847 | 549 |
| | TMA | $4.7 \times 10^{8}$ | 436 | 614 | $1.0 \times 10^{7}$ | 10307 | 1030 |
| | Cho | - | - | - | $1.19 \times 10^{7}$ | 11213 | 9440 |

See SI Table 2 for details of background (bkg) constructs for Nic and Ana (bkg: βL262S + δL265S + δP123R). See Nayak et al., 2017 for the details of the rest of the agonist binding kinetics at all the three binding sites (17).

SI Table 2. Rates and equilibrium constants for mutant constructs of fetal AChRs.

| Mutant constructs | $f_0/f_2'$ | $b_0/b_2'$ | $E_0/E_2'$ |
|---|---|---|---|
| βL262S + δL265S + δP123R | 99 | 7777 | 0.013 |
| βL262S + γL260Q + δP123R | 109 | 2333 | 0.047 |
| αY198F + βL262S + δL265S + δP123R | 83 | 5893 | 0.014 |
| αA96H + βV266A | 1443 | 85 | 16.97 |
| βL262S + δL265S + δP123R + 100 pM ACh | 89 | 5056 | 0.018 |
| βL262S + δL265S + δP123R + 100 pM ACh + 20% glycerol | 21 | 6497 | 0.0032 |

Single channel kinetic rate constants ($f_0/f_2'$ and $b_0/b_2'$), equilibrium constants ($E_0=f_0/b_0$ or $E_2=f_2'/b_2'$). The rate and equilibrium constants for the mutant constructs where [ACh] is not mentioned are $f_0$, $b_0$, and $E_0$.

SI Table 3. MD simulation data for understanding binding site dynamics of the WT vs mutant AChRs.

| Structure Parameters | WT (Average±SD) | Mutant (major component) (Average±SD) | Mutant (minor component) (Average±SD) |
|---|---|---|---|
| Volume: α–γ site (Å$^3$) | 133.6±11.19 | 173.1±18.59 | 199.9±5.32 |
| Volume: α–δ site (Å$^3$) | 139.7±4.09 | 137.7±4.45 | - |
| Distance: αY190 (Cα) - CCh (QA) (Å) | 6.1±0.94 | 11.7±2.37 | 15.2±1.35 |
| Distance: αC192 (Cα)-CCh (QA) (Å) | 12±1.45 | 17±2.45 | 21.5±1.15 |
| Distance: αW149 (Cα)-αC192 (Cα) (Å) | 16.4±1.33 | 21.9±1.99 | 24.6±1.89 |
| Distance: αK145 (Cα)-αD200 (Cα) (Å) | 3.1±1.06 | 4.8±0.82 | - |
| Dihedral: CCh-τ1 (˚) | +63±17.07/ -62±15.91 | +62±17.18/ -65±15.46 | - |
| Dihedral: γW55-χ1 (˚) | +51±8.64 | -50±10.4 | - |

SI Table 4. Rates and equilibrium constants for AChRs with charge neutralization mutations (γE57A+γD113A+γD174A+γE180A).

| $f_0$ ($s^{-1}$) | $b_0$ ($s^{-1}$) | $E_0^{obs}$ | $j_{on}$ ($M^{-1}.s^{-1}$) | $j_{off}$ ($s^{-1}$) | $J_d$ (nM) |
|---|---|---|---|---|---|
| 119 | 5993 | 0.2 | $5.1 \times 10^9$ | 197 | 38.6 |
| $f_1$ ($s^{-1}$) | $b_1$ ($s^{-1}$) | $E_1^{obs}$ | $k_{on}$ ($M^{-1}.s^{-1}$) | $j_{off}$ ($s^{-1}$) | $K_d$ (μM) |
| 2119 | 113 | 18.8 | $4.3 \times 10^7$ | 3721 | 86.5 |

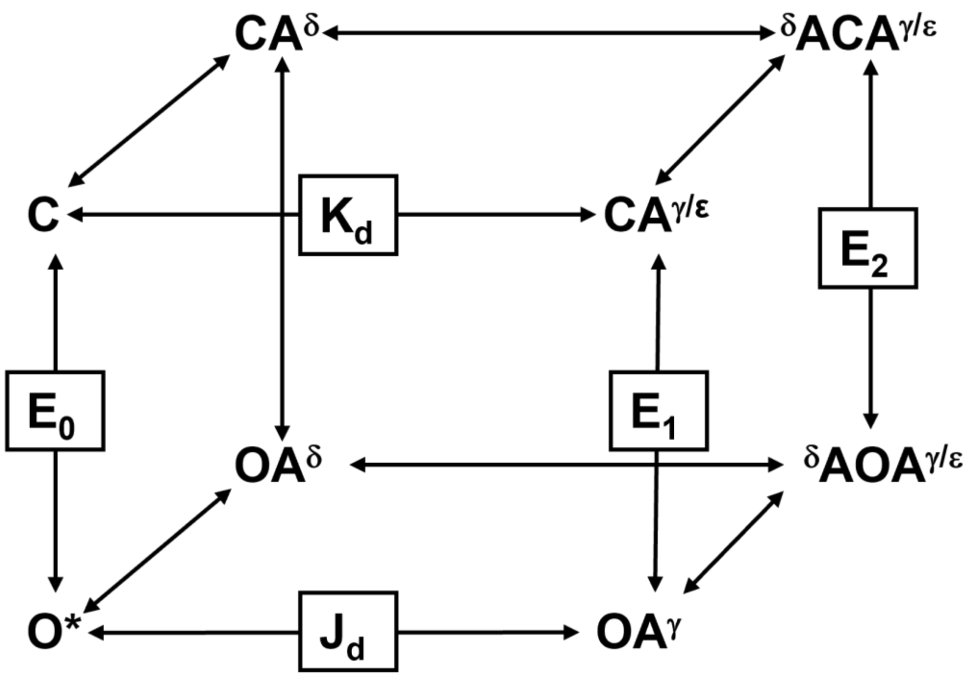


**SI Figure 1. Cube kinetic model of AChR activation.** C, O, O*, A, CA, and OA have the usual meaning (see SI Methods). Vertical and horizontal arms represent gating and binding steps. $E_n$ = gating equilibrium constants (n = 0-2 where n represents the number of bound agonists). $K_d$, $J_d$= low- and high-affinity dissociation constants.

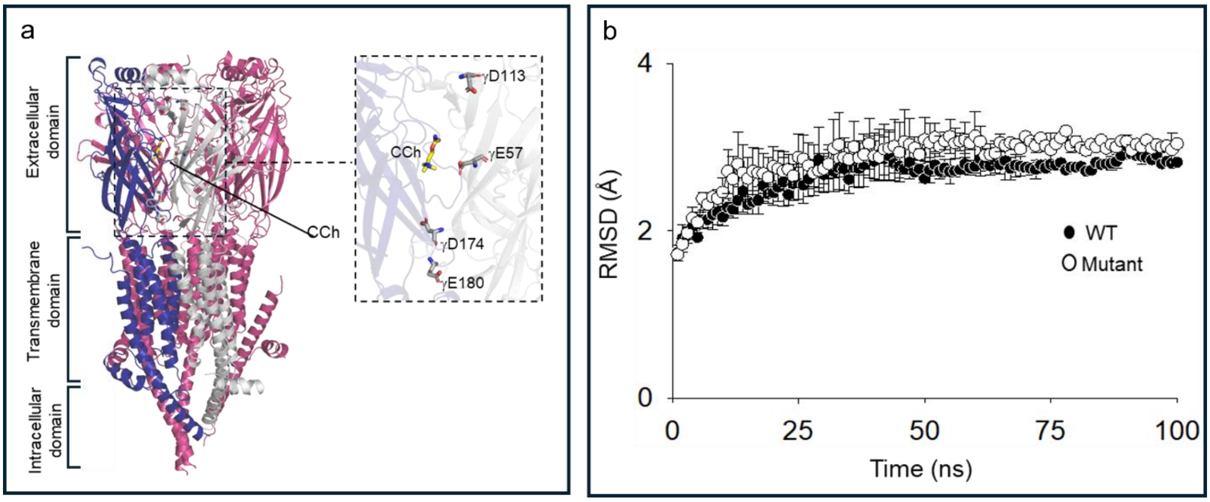


**SI Figure 2. Structure of AChR and stability of MD simulations.** a. Membrane view of the AChR (PDB Id:7QL6) bound with the ligand carbamylcholine (CCh; yellow) (left). On the right see an enlarged view of the receptor showing the charged residues (γE57, γD113, γD174, γE180) that are mutated to alanine for the present study. b. Line and scatter plot of $C_\alpha$ RMSD (mean±SD, triplicates) of 100 ns of all atom MD simulation run. The filled and open circles represent WT and mutant runs. The structure stabilizes at ~3Å in both WT and mutant simulation runs.

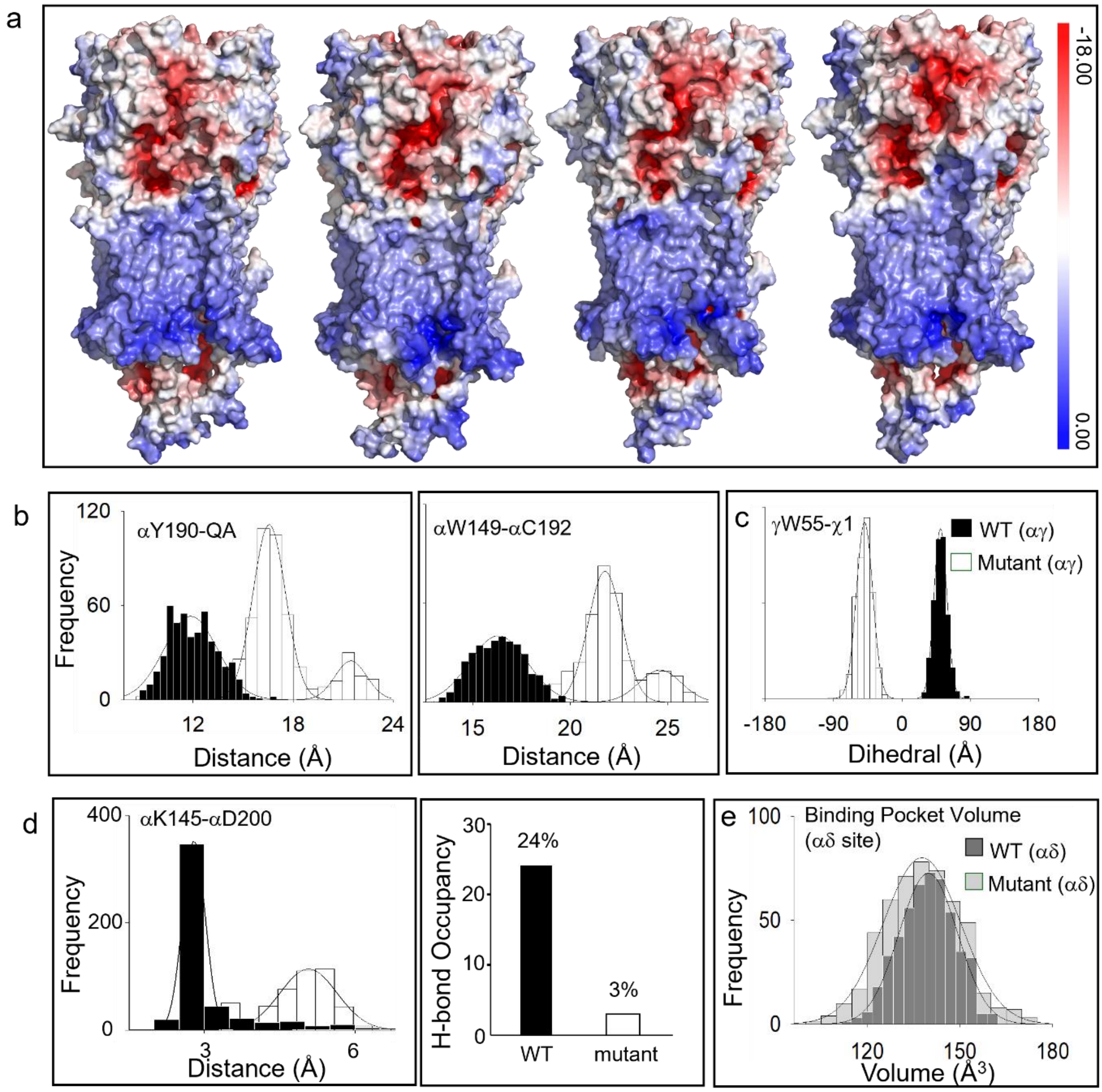


**SI Figure 3. Effect of reduction in negative charges in ECD of the γ subunit on the overall structural dynamics**. **a**. Electrostatic potential map of the AChRs (PDB Id:7QL6) with single mutation γE57A, γD113A, γD174A, and γE180A (left to right). Single residue mutation does not affect the electrostatic potential significantly. **b and c**. Histograms showing the distance between αY190-LIG(QA) (left), and αW149-αC192 (middle) and dihedral angle distribution of γW55-χ1 (right). Solid spline curve: Gaussian fit of the histogram peaks. Mean and standard deviation values of the fit are in SI Table 3. **d.** Histogram showing the comparison between the αK145-αD200 salt bridge of WT and mutant receptor (left). The average distance between αK145-αD200 in WT and mutant receptor is 3.1±1.06 Å and 4.8±0.82 Å (SI Table 3). The bar graph on the right shows the percentage of H-bond occupancy between αK145 and αD200 in the WT and mutant receptor. **e**. Histogram showing the overlay of volume distribution at the α−δ interface (αY93, αW149, αY190, and δW57) for the WT (average volume = 139.7±4.09 $Å^3$) vs the mutant (average volume = 137.7±4.45 $Å^3$) receptor.
